# Predicting the presence and abundance of bacterial taxa in environmental communities through flow cytometric fingerprinting

**DOI:** 10.1101/2021.05.05.442872

**Authors:** Jasmine Heyse, Florian Schattenberg, Peter Rubbens, Susann Müller, Willem Waegeman, Nico Boon, Ruben Props

**Author notes:** Correspondence to: Nico Boon, Ghent University; Faculty of Bioscience Engineering; Centre of Microbial Ecology and Technology (CMET); Coupure Links 653; B-9000 Gent, Belgium; Webpage: www.cmet.ugent.be.

## Abstract

Microbiome management research and applications rely on temporally-resolved measurements of community composition. Current technologies to assess community composition either make use of cultivation or sequencing of genomic material, which can become time consuming and/or laborious in case high-throughput measurements are required. Here, using data from a shrimp hatchery as an economically relevant case study, we combined 16S rRNA gene amplicon sequencing and flow cytometry data to develop a computational workflow that allows the prediction of taxon abundances based on flow cytometry measurements. The first stage of our pipeline consists of a classifier to predict the presence or absence of the taxon of interest, with yields an average accuracy of 88.13±4.78 % across the top 50 OTUs of our dataset. In the second stage, this classifier was combined with a regression model to predict the relative abundances of the taxon of interest, which yields an average R^2^ of 0.35±0.24 across the top 50 OTUs of our dataset. Application of the models on flow cytometry time series data showed that the generated models can predict the temporal dynamics of a large fraction of the investigated taxa. Using cell-sorting we validated that the model correctly associates taxa to regions in the cytometric fingerprint where they are detected using 16S rRNA gene amplicon sequencing. Finally, we applied the approach of our pipeline on two other datasets of microbial ecosystems. This pipeline represents an addition to the expanding toolbox for flow cytometry-based monitoring of bacterial communities and complements the current plating- and marker gene-based methods.

**Importance:** Monitoring of microbial community composition is crucial for both microbiome management research and applications. Existing technologies, such as plating and amplicon sequencing, can become laborious and expensive when high-throughput measurements are required. Over the recent years, flow cytometry-based measurements of community diversity have been shown to correlate well to those derived from 16S rRNA gene amplicon sequencing in several aquatic ecosystems, suggesting there is a link between the taxonomic community composition and phenotypic properties as derived through flow cytometry. Here, we further integrated 16S rRNA gene amplicon sequencing and flow cytometry survey data in order to construct models that enable the prediction of both the presence and the abundance of individual bacterial taxa in mixed communities using flow cytometric fingerprinting. The developed pipeline holds great potential to be integrated in routine monitoring schemes and early warning systems for biotechnological applications.

## Introduction

Bacterial communities are complex and highly dynamic associations that play important roles in many biotechnological applications. One issue that hinders efforts to study and manage these communities, is the fact that existing technologies to assess community composition either rely on cultivation or necessitate the extraction and sequencing of genomic material, both of which are time consuming and laborious. As a result, the availability of fine-scale resolution data on bacterial community dynamics is still limited in many fields. One example hereof is the aquaculture sector (Wang *et al*., 2020), where the development of effective management strategies to reduce the occurrence of diseases is hampered by the limited knowledge on the microbial ecology of these systems. Additionally, routine monitoring schemes in aquaculture farms are still mainly relying on (selective) plating, which prohibits accurate description of general dysbiotic states and specific disease outbreaks.

Flow cytometry (FCM) is a single-cell technique that is increasingly used as a fast and inexpensive tool for characterising microbial communities in a wide variety of fields, including drinking water production and distribution (Besmer and Hammes, 2016; Buysschaert, Vermijs, *et al*., 2018; Favere *et al*., 2020), surveys of natural ecosystems (Ferrera *et al*., 2015; Read *et al*., 2015; Santos *et al*., 2019; Giljan *et al*., 2020), aquaculture (Lucas *et al*., 2010) and fermentation (Salma *et al*., 2013; Narayana *et al*., 2020). Over the last decade, through the development of advanced data-analysis pipelines, the application of FCM has moved beyond its initial purpose of estimating cell densities (Rubbens and Props, 2021). These computational advances include a range of fingerprinting pipelines (Koch, Fetzer, Harms, and Muller, 2013; Koch, Fetzer, Schmidt, *et al*., 2013), algorithms for estimating community stability (Liu *et al*., 2018) and algorithms for estimating community diversity metrics (Props *et al*., 2016). Flow cytometry-derived diversity metrics have been shown to be highly correlated to those derived from 16S rRNA gene amplicon sequencing in some ecosystems (García *et al*., 2015; Props *et al*., 2016, 2018; Rubbens *et al*., 2021), suggesting there is a link between the taxonomic community composition and phenotypic properties as derived through FCM. This observation is supported by the fact that sorted fractions of a community have different taxonomic compositions compared to the entire community (Vogt *et al*., 2009; Zimmermann *et al*., 2016; Lambrecht *et al*., 2019; Liu *et al*., 2019; Haange *et al*., 2020).

Using machine learning techniques, Bowman *et al*. (2017) and Rubbens, Schmidt, *et al*. (2019) showed that the relative abundance of specific OTUs is predictive for the abundance of high nucleic acid (HNA) and low nucleic acid (LNA) sub-communities in FCM data of natural ecosystems, illustrating the possibility of linking specific regions in the cytometric fingerprint to taxonomic groups using modelling approaches. Several studies have sought to further exploit this relationship in order to build predictive models for taxonomic community composition based on FCM data. Most of these studies take a bottom-up approach in which they train predictive models on data of axenic bacterial cultures. Rubbens *et al*. (2017) introduced the use of *in silico* communities based on axenic culture data, while Özel Duygan *et al*. (2020) developed a pipeline that allows to classify mixed communities into classes of predefined “cell types” by comparing data to signatures of a set of strains and bead standards. However, cytometric fingerprints of axenic cultures are known to be dynamic over time, for example in function of growth stage (Müller, 2007; Neumeyer *et al*., 2012; Buysschaert, Kerckhof, *et al*., 2018). Additionally, we have recently shown that the single-cell properties of an individual taxon, as measured by FCM, depend on the presence of other bacterial taxa in the community. Therefore, training models on axenic culture data may lead to unreliable predictions (Heyse *et al*., 2019).

In this study, we aimed to further integrate 16S rRNA gene amplicon sequencing and flow cytometry survey data in order to construct models that enable the prediction of both the presence and the abundance of multiple individual bacterial taxa in mixed communities using flow cytometric fingerprinting (Figure 1). As a case study, we used samples taken from a whiteleg shrimp (*Litopenaeus vannamei*) hatchery of which the dynamics have been previously described (Heyse *et al*., 2021). We first verified the taxonomic stratification in the cytometric fingerprints using cell sorting. We then developed a two-stage pipeline using flow cytometry data as input that, firstly, predicts the presence/absence of bacterial taxa, and, secondly, predicts the relative abundance of bacterial taxa. Through the direct linking of flow cytometry and amplicon sequencing survey data, the constructed models are not relying on data from axenic cultures. We verified the ability of the models to assign taxa to the specific regions in the cytometric fingerprint using marker gene data from the cell sorted community fractions and using a three strain mock community. Finally, we validated the approach of our pipeline on two independent datasets.

**Figure 1.**
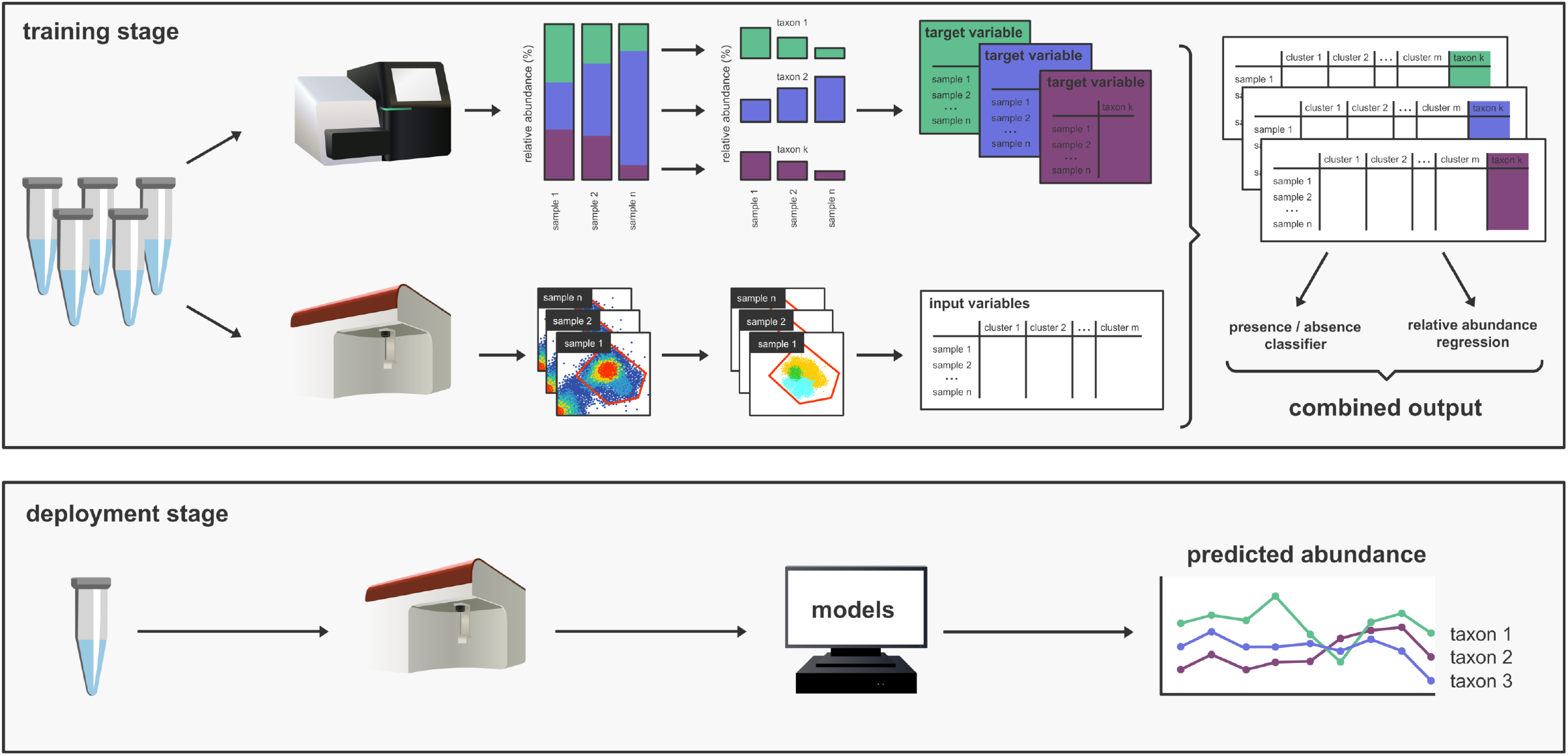
Overview illustration of the workflow and application of the pipeline presented in this study. During the training stage, samples from the system under study are collected and analysed using both flow cytometry and 16S rRNA gene amplicon sequencing. For the 16S rRNA gene amplicon data, the reads are processed to calculate relative abundance profiles for each sample. The models are trained for each taxon individually. Therefore, the relative abundances of the taxa of interest are extracted which results in a single vector for each taxon. For flow cytometry, the single cell data are separated from the background signals by manually creating a gate on the primary fluorescent channels and subsequently discretised by applying a Gaussian Mixture mask, which assigns each cell to a specific cluster. This results in a data frame with the relative abundance for each cluster of the Gaussian Mixture in each sample. Two models are constructed for each taxon: an absence/presence classifier and a regression ensemble to predict the relative abundance of the taxon of interest. During the deployment stage, the system under study is sampled using flow cytometry, the trained models are used to predict the presence/absence and relative abundances of one or multiple taxa of interest.

## Results

In this study, we used published flow cytometry and 16S rRNA gene amplicon data from an 18-day sampling campaign in a *L. vannamei* hatchery where five replicate cultivations were studied (Heyse *et al*., 2021). The replicate cultivation tanks were sampled at a resolution of 3 hours for flow cytometry and once per day for 16S rRNA gene sequencing. This dataset was combined with newly-generated 16S rRNA gene amplicon data on sorted fractions of samples originating from this previous study.

### Taxonomic information is conserved in flow cytometric fingerprints

Prior to model training, the connection between the taxonomic composition of the bacterial communities, as derived through 16S rRNA gene amplicon sequencing, and their phenotypic properties, as derived by flow cytometry, was evaluated using cell sorting. In total, 57 community fractions were sorted from 20 samples using 5 gates (referred to as “sub-community” or “SC” 1 to 5). The sorted regions in the flow cytometry data space (i.e. gates) were chosen to maximize the coverage of the community across the side scatter and SYBR Green I fluorescence range (Supplementary Figure 1), and represented sub-communities with relative cell abundances between 3 to 56 % of the total cell gate (Figure 2A).

**Figure 2.**
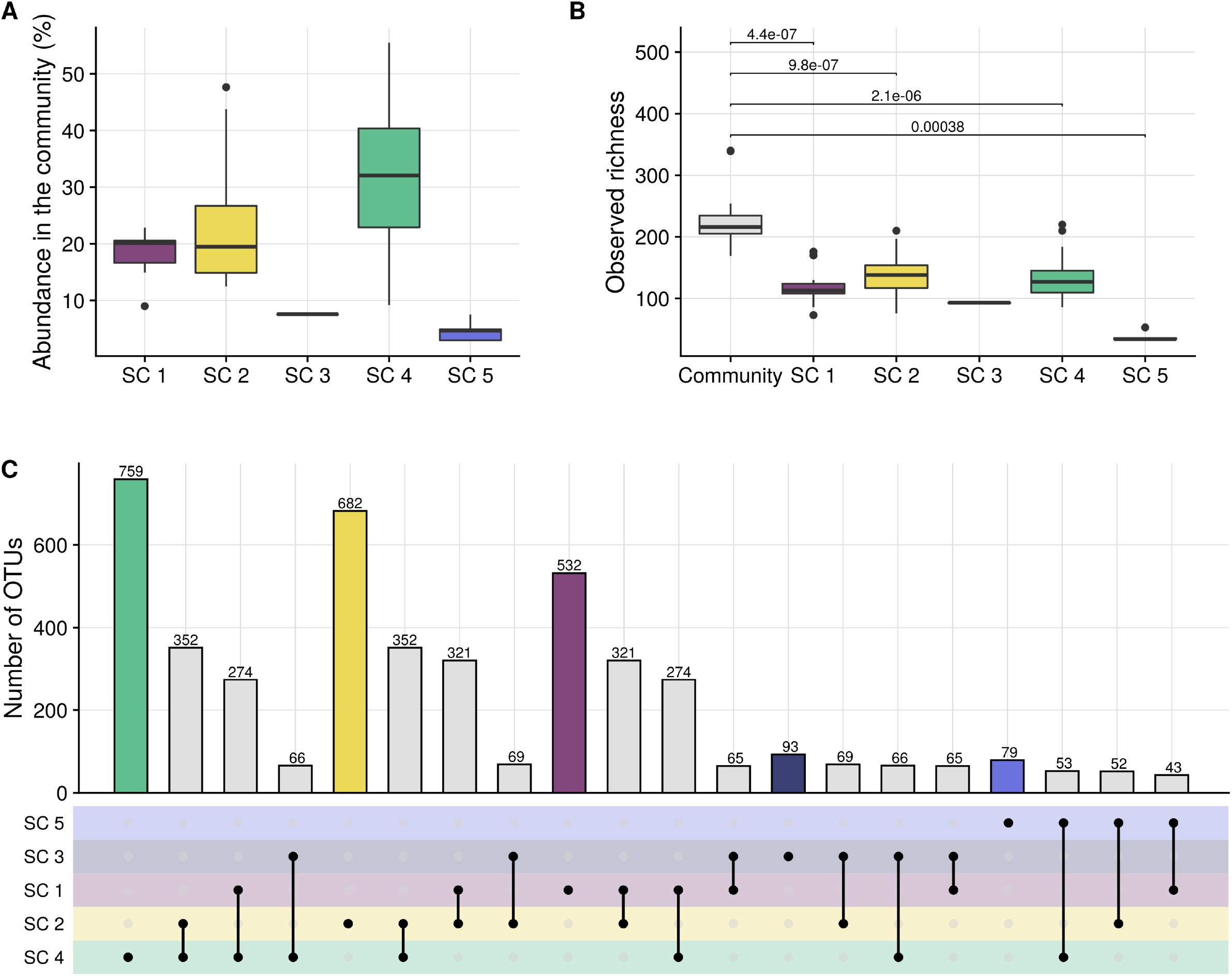
(A) Relative abundances of the sorted sub-communities (SC), based on the measurements of the Influx v7 Sorter. (B) Observed taxonomic richness in the sorted community and sub-communities. The values above the brackets indicate the p-values of a one-sided (lower) Wilcoxon rank sum test. Note that for sub-community 3 no p-value is supplied since this sub-community was sorted only once. (C) Upset graph illustrating intersections between the taxonomic composition of the sorted sub-communities (i.e. number of common OTUs). The upper bars illustrate the cumulative number of OTUs that are found in a sub-community (in case of a single dot) or shared between sub-communities (in case of two connected dots). Note that the number of sorted samples were not homogeneously distributed over the five sorting gates (i.e. SC3 and 5 were sorted once and three times, respectively, while SC1, 2 and 4 were sorted 15, 17 and 18 times, Supplementary Figure 2).

For all sub-communities, the taxonomic richness was significantly lower as compared to that of the cell gate (one-sided Wilcoxon rank sum test, p < 0.0001, Figure 2B). The taxonomic composition of each of the five gated sub-communities was significantly different from that of the cell gate as well as from each other (PERMANOVA on Bray-Curtis dissimilarities, p < 0.01, Supplementary Table 1, Supplementary Figure 1). Each sub-community was enriched in specific taxa and shared a limited number of taxa with the other sub-communities (Figure 2C). Many taxa were uniquely detected in a specific sub-community (e.g. OTU1 *Phaeodactylibacter* sp. in SC 1), however, some taxa were detected in two (e.g. OTU3 *Nioella* sp. in SC 1 and 2) or three (e.g. OTU7 *Kordia* sp. in SC 1, 2 and 3) sub-communities (Figure 2C, Supplementary Figure 2). The overlap in taxonomic composition between gates that were more dissimilar from each other was smaller (e.g. SC 1 and 5, which are more dissimilar, only share 15 OTUs, while SC 1 and 2, which are close to each other, have 147 OTUs in common; Figure 2C), confirming that specific taxa typically occur in the specific positions of the cytometric space.

The two most narrowly defined sub-communities (i.e. SC 3 and 5), with the lowest abundance in the community, represented sub-communities with low taxonomic diversity and were nearly mono-dominant, (i.e. *Kordia* sp. in SC 3 and unclassified *Alphaproteobacteria* sp. in SC 5), while the larger and abundant gates (i.e. SC 1, 2 and 4) were dominated by multiple taxa (Supplementary Figure 2). It should be noted that the number of sorted samples were not equally distributed over the five sorting gates (i.e. SC3 and 5 was sorted once and three times, respectively, while SC1, 2 and 4 were sorted 15, 17 and 18 times), which may have caused the cumulative number of observed taxa in SC3 and 5 to be lower than those of SC1, 2 and 4. Nevertheless, also the average number of taxa per sample was lower in SC3 and 5 as compared to SC1, 2 and 4 (Figure 2B).

Throughout the shrimp cultivation, the phylogenetic composition in the sub-communities was preserved well, even though the composition of the total community was dynamic over time and differed between the replicate tanks from which samples were sorted.

### Development of a pipeline to extract taxonomic information

Cell sorting was performed on a different instrument (BD Influx) as compared to the FCM measurements of community samples (BD FACSVerse). To be able to use both the community sample and the sorted sample data as a single dataset, a set of representative samples was measured on both instruments, the gates that were used for sorting were manually-recreated on the FACSVerse data and correspondence between the relative cell abundances in the gates on data of the two instruments was used to evaluate the quality of the manually recreated gates (Supplementary Figure 1). The corresponding flow cytometric fingerprints of the sorted sub-communities were obtained from the community measurements using these gates. The combined dataset (i.e. including both sorted and community measurements) consisted of 169 samples for which both 16S rRNA gene amplicon and flow cytometry data were available. Models were trained for each OTU individually, using the flow cytometry data as input and the presence or abundance of the OTU of interest as model output. Details about the model construction are provided in the Materials and Methods sections. Performances for the top 50 OTUs from the aquaculture dataset were evaluated. All reported performance values are performances on the validation sets (i.e. on data that was not used for model training).

In the first part of the pipeline, a presence/absence classifier is trained. Classification performance was evaluated using accuracy (i.e. percentage correctly predicted samples) and AUC (area under the ROC curve, i.e. probability that a randomly-chosen sample where the taxon is “present” is assigned a higher probability for “present” than a randomly-chosen sample where the taxon is “absent”). We were able to perform presence/absence classification with high accuracies, ranging from 78 % to 98 % for individual OTUs and AUC values between 0.66 and 0.99 (Figure 3A and B). The number of false positive (i.e. taxon is incorrectly predicted to be present) and false negative (i.e. taxon is incorrectly predicted to be absent) samples did not differ strongly for individual OTUs (two-sided Wilcoxon rank sum test, p > 0.05, Supplementary Figure 3).

**Figure 3.**
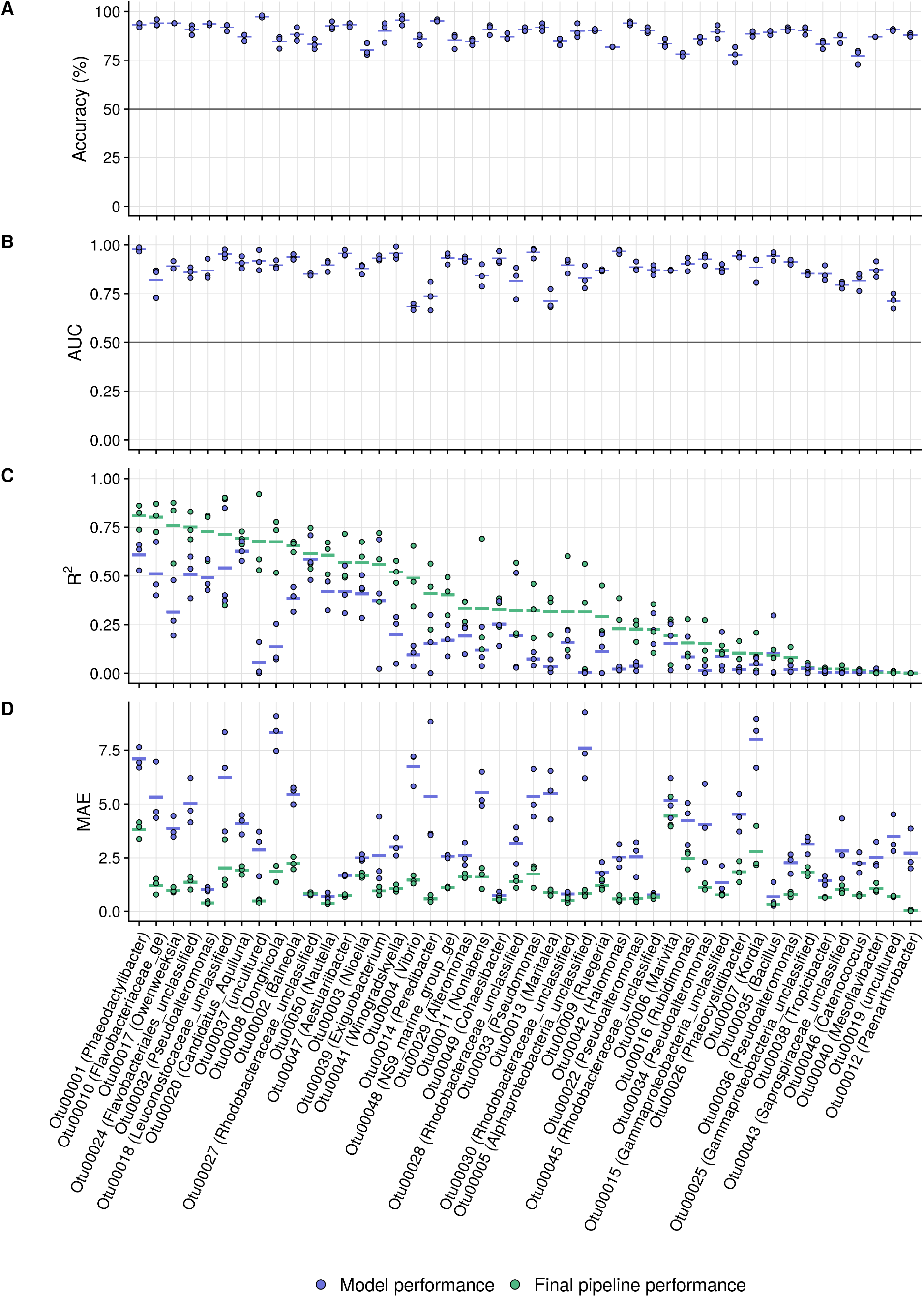
Classifier accuracy (A) and AUC (B), and regression R^2^ (C) and MAE (D) values for the top 50 abundant OTUs from the aquaculture dataset. For the regression metrics (R^2^ and MAE) both the regression model outputs (in blue) and final pipeline outputs (i.e. after imposing the classifier predictions to the regression results, in green, visualised in Supplementary Figure 4) are illustrated. OTUs are ordered according to their final R^2^ values. The three dots for each model represent three repeated fold splits, the vertical line per OTU indicates the average performance of the replicates. The vertical line at 50 % in (A) and 0.5 in (B) indicates the random guessing threshold of a binary classifier.

In the second part of the pipeline, the relative abundances of individual taxa were modelled using a regression ensemble. Regression performance was evaluated using R^2^ (i.e. proportion of the variance in the relative abundance values that can be predicted from the flow cytometry data) and MAE (mean average error, i.e. average deviation between true and predicted relative abundances).The regression ensembles had R^2^-values between 0.00 and 0.64 (0.21 ± 0.18 on average) and MAE (Mean Absolute Error) values between 0.24 and 9.06 (3.41 ± 2.19 on average) (blue dots in Figure 3). The regression ensembles frequently predicted high relative OTU abundances for samples where an OTU was either absent or present in very low abundance (Supplementary Figure 4B). Therefore the predictions of the classifier were superimposed on the regression predictions (Supplementary Figure 4A): the predicted relative OTU abundances in samples that were classified as “absent” were set to zero, predictions for samples where the OTU was predicted to be “present” remain unchanged. This reduced the number of false positive samples by an average of 10 fold (i.e. from 40 ± 17 to 4 ± 3 out of 100 samples). However, superimposing the classifier to the regression results slightly increased the number of false negative samples from 3 ± 3 out of 100 samples to 8 ± 5 on average. Overall, the R^2^-values were increased to 0.35 ± 0.24 on average (ranging between 0.00 and 0.81), and the MAE was reduced to 1.31 ± 0.97 on average (green dots in Figure 3).

To evaluate the ability of our approach to correctly capture dynamics of taxa over time, we predicted the presence and relative abundances of four taxa on the time points for which no amplicon data were available. Additionally, we calculated the predicted absolute OTU abundances by multiplying bacterial densities with the predicted relative OTU abundances. The taxa were selected based on a good (OTU1, R^2^ = 0.81), intermediate (OTU2, R^2^ = 0.65 and OTU6, R^2^ = 0.19) and low (OTU13, R^2^ = 0.03) overall prediction performance. For OTU1, the predictions followed the overall patterns that were estimated by interpolation of the time points for which amplicon data was available (Figure 4). Additionally, the predictions for which the abundances did not match the trends that were estimated by interpolation, often coincided with low absolute abundances. Similarly, for OTU2 and OTU6, which had intermediate model performances, the abundance patterns were following the expected trends well (Supplementary Figure 5, Supplementary Figure 6). For OTU13, which had the lowest performance, the patterns were not corresponding to those that would be expected based on interpolation of the available data points (Supplementary Figure 7).

**Figure 4.**
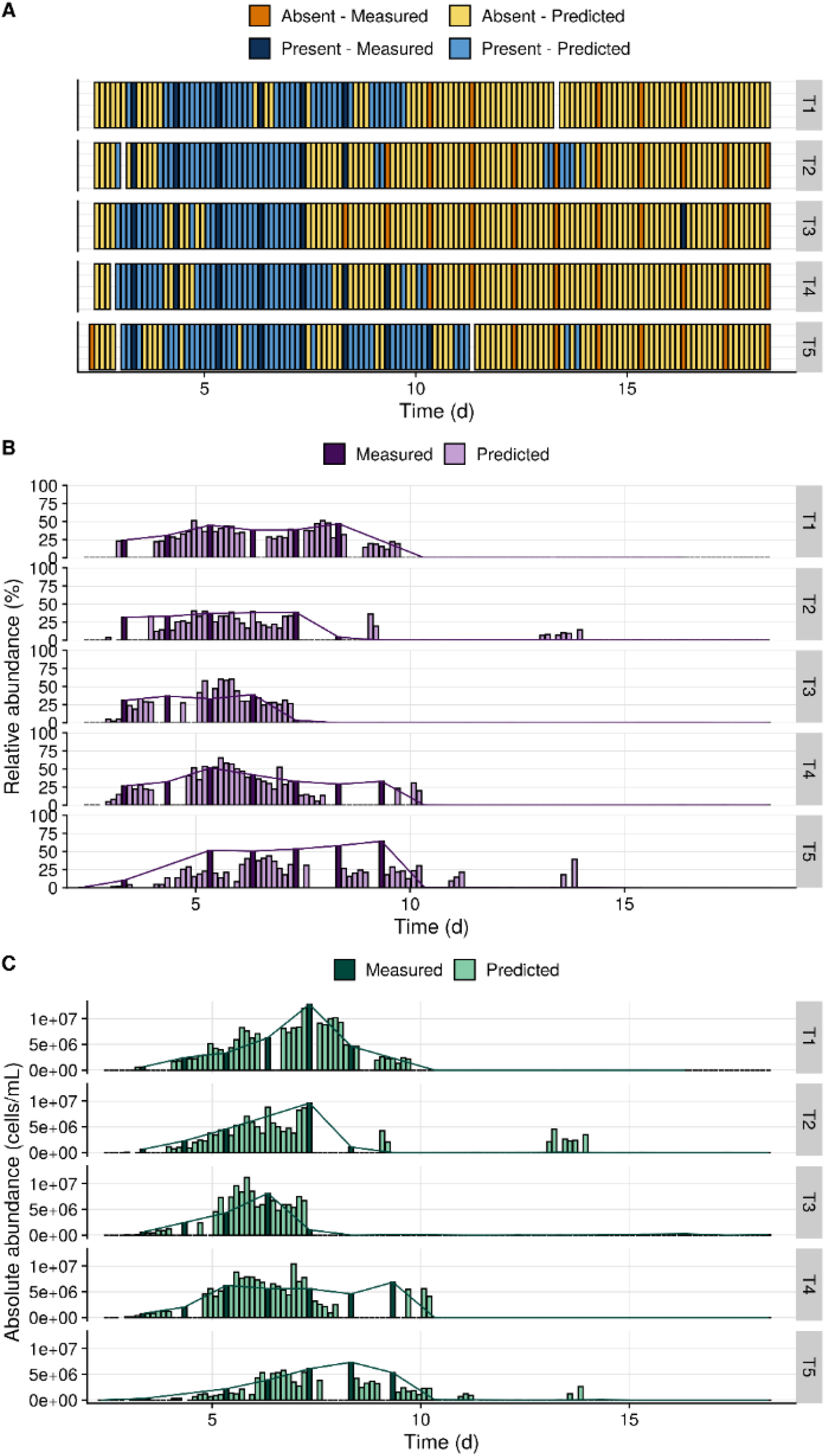
Predictions for OTU 1 (*Phaeodactylibacter* sp.; R^2^ = 0.81) from the aquaculture dataset. The five replicate shrimp cultivation tanks (“T1” to “T5”) were sampled at a resolution of 3 hours for flow cytometry and once per day for 16S rRNA gene sequencing. The presence and relative abundances for OTU1 on the time points for which no amplicon data were available were predicted in order to evaluate the ability of our approach to correctly capture dynamics of this taxon over time. The dark shades (“measured”) correspond to the values that were determined based on 16S rRNA sequencing. The lighter shades (“predicted”) correspond to time points for which only flow cytometry data was available and predictions were made using the models. Expected values can be estimated by interpolation of the measured samples (indicated with the lines between the measured samples). The reported values are averages of the two replicate measurements at each time point. (A) Predictions of the presence/absence classifier. (B) Predicted relative abundances. (C) Predicted absolute abundances, calculated by multiplying the predicted relative abundances by the total cell density as determined through flow cytometry.

Since the models were trained on survey data, in which there may be co-occurrence between taxa, predictions of individual OTUs may be (partly) relying on detecting co-occurring OTUs and not the OTU of interest itself. In that case, the applicability of the pipeline may be limited to filling gaps in time series of the dataset that was used for model training (i.e. relying on auto-correlation between the samples over time), but the reliability of predictions on independently generated time series of the same environment (e.g. repeated shrimp cultivation in this case) may be limited. To verify the impact of co-occurrence, we compared the performances of models that were trained on only four of the replicate tanks and predictions were made on the 5^th^ tank (setting 1) with models that were trained using a randomly chosen training- and validation set from data of all replicate tanks (setting 2). The former ensured that the co-occurrence patterns of the validation data (i.e. data from the 5^th^ tank) were not incorporated during model training, while the latter incorporated a all co-occurrence patterns during model training. There was an average decrease in R^2^ of 0.02 across the 50 OTUs in setting 2 relative to setting 1. This small decrease suggests that co-occurrence has only a minor influence on model performance. To investigate this further, we assessed, for the top 10 OTUs, the feature importance of the clusters in the cytometric fingerprint (see Materials and Methods for procedure) with the regions of the sorting gates in which these taxa were observed. Overall, the positions of clusters with high feature importances were corresponding well to the positions of the gates in which these taxa were observed, with the exception of OTU6, for which clusters were detected over the entire range of the bacterial community fingerprint (Supplementary Figure 8). For some OTUs there were small deviations, which may be the result of technical aspects. For example, some OTUs were not detected in regions with high feature importances, which may be the result of the limited number of sorted samples and the fact that these were biased towards only 3 tanks during the first half of the sampling campaign (i.e. day 4-13). Secondly, the sorting gates were recreated from the data of one instrument to the other (see Materials and Methods, Supplementary Figure 1). This may have caused gates immediately adjacent to the sub-communities to be either marked or not marked while this was not the case. Overall, these results show that the models can robustly associate taxa to regions in the cytometric fingerprint where they are detected using 16S rRNA gene amplicon sequencing, and, hence they are not relying heavily on co-occurrence patterns.

To test whether taxa that are phylogenetically closely related are more likely to be associated to the same regions in the cytometric fingerprints, the relationship between phylogenetic distance between taxa and feature importance similarity was evaluated. There was a significant (Adj. R. sq. = 0.039 and p < 2e-16, C_p_ = −0.20) relationship between the similarity of cluster importance for different OTUs assigned by the model and the phylogenetic similarities (Supplementary Figure 9). This relationship was negative, indicating that OTUs which are phylogenetically more closely related, are more likely to be associated with the same regions in the cytometric fingerprints.

The sensitivity of the model performance to the amount of data available for training was investigated for two OTUs (i.e. OTU1 and 6), by training models on randomly subsampled datasets that contained 20, 40, 60 or 80 % of the dataset (i.e. 34, 68, 101 or 135 samples). For both OTUs and for both classification and regression, there was a strong reduction in performance at the lower sample sizes (learning curves in Supplementary Figure 10). Classification accuracy was reduced by 10 % and 5 %, for OTU1 and OTU6, respectively, for every 20 % reduction in dataset size. For the regression models, the R^2^-values were halved when the model was trained on only 20 % of the data as compared to when it was trained on 80 % of the data. For both of the OTUs the performance did not yet reach a plateau, suggesting that more data is required to improve model performances.

### Validation of the approach on external datasets

To test whether the approach of our pipeline was applicable for monitoring of other (managed) microbial systems, the entire workflow was replicated on a three strain cytometric mock community from Cichocki *et al*. (2020) and a dataset of insular reactor communities from Liu *et al*., (2019). Details about the datasets are provided in Supplementary Table 2.

For the mock community classifier AUC was 0.96 ± 0.07 % on average and R^2^-values were 0.89 ± 0.03 on average (Figure 5). Since this was a simple mock community, we could validate that the clusters that were assigned a high importance by the model corresponded well to the regions where these taxa were found in the cytometric fingerprint (Supplementary Figure 11). For the reactor communities, AUC of the top 18 OTUs were 0.81 ± 0.12 on average. As for the aquaculture dataset, there were big differences in the model performances of individual OTUs. The range of performances was similar as for the aquaculture dataset, with an average R^2^ of 0.33 ± 0.27.

**Figure 5.**
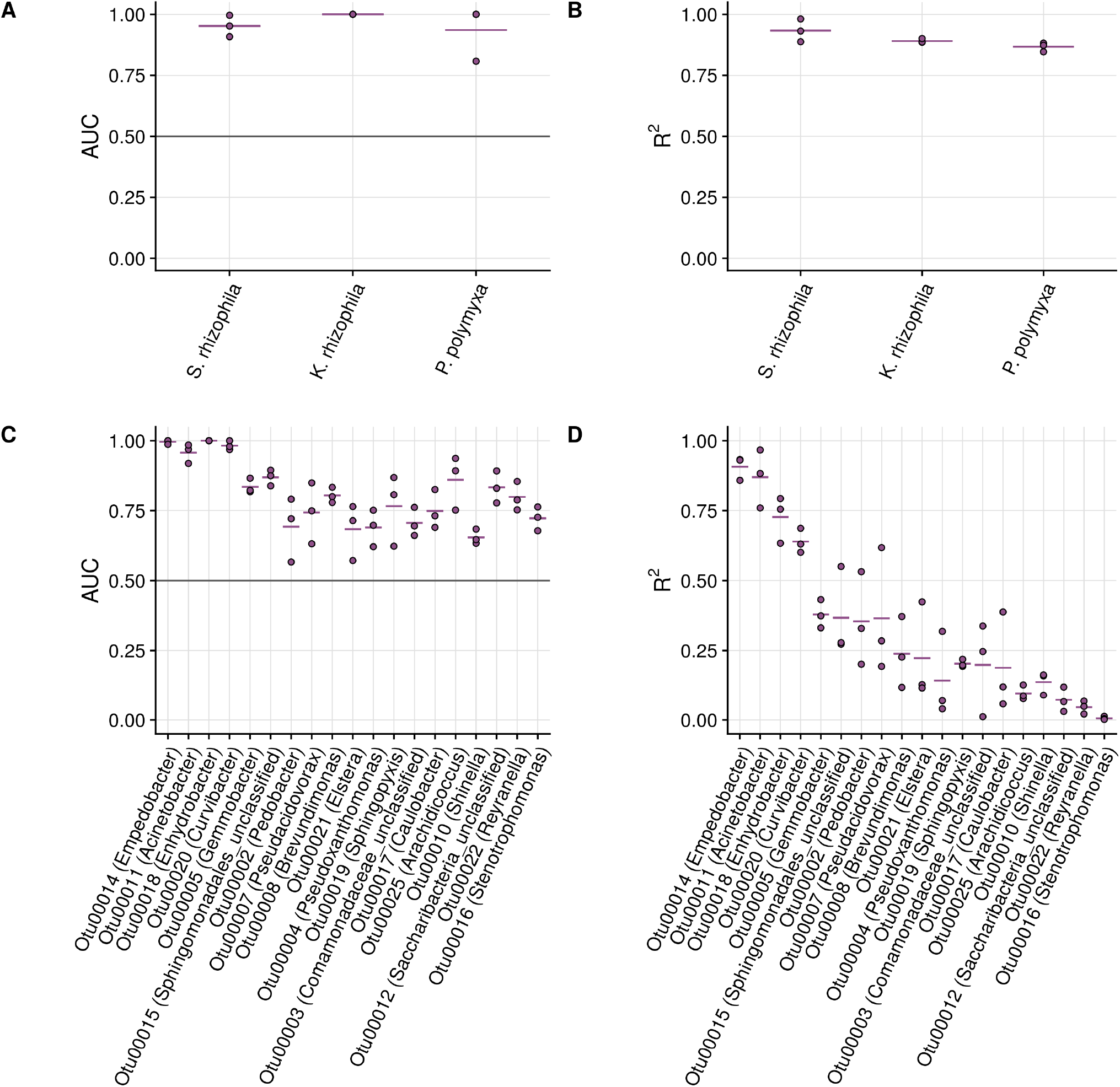
Model performances on the two validation sets. (A) Classifier AUC-values for the three strain mock community. (B) R^2^ values for the three strain mock community. (C) Classifier AUC-values for the top 18 OTUs of the reactor communities. (D) R^2^ values for the top 18 OTUs of the reactor communities. The three dots for each model represent three repeated fold splits, the vertical line per OTU indicates the average performance of the replicates. The vertical line at 0.5 in (A) and (C) indicate the random guessing threshold of a binary classifier.

## Discussion

The objectives of our study were: (1) to verify the taxonomic structure in flow cytometry fingerprints for our model system, *L. vannamei* larviculture rearing water communities, using cell sorting; (2) to further integrate 16S rRNA gene amplicon sequencing and flow cytometry data to develop a pipeline that allows to predict the presence/absence and the relative abundance of multiple individual bacterial taxa in mixed communities based on flow cytometry measurements (Figure 1); (3) to validate the approach of our pipeline on two independent datasets.

### Models can predict temporal abundance dynamics

Substantial variation in model performances were observed for the individual OTUs, for both the aquaculture (Figure 3) and the validation datasets (Figure 5). For all OTUs the classifier accuracies were largely above the random guessing threshold of 50 %, indicating that the presence of all taxa could be predicted with moderate to high accuracy. In contrast, for the prediction of relative abundances, there were large differences in performance between OTUs. For the aquaculture dataset, predictions for OTUs with a high to intermediate R^2^ occasionally diverged from what would be expected based on interpolation of the time points for which 16S rRNA data was available, but the overall patterns of taxon presence and abundance were predicted well (Figure 4, Supplementary Figure 5, Supplementary Figure 6). Based on these results we conclude that the constructed models are suitable for monitoring dynamics over time, but that one should be more cautious when evaluating single snapshot samples. The number of required samples to predict reliable trends will be dependent on the taxa of interest and the dynamics of the system under study. We acknowledge that for a subset of the investigated OTUs, accuracies were very low and predictions were not corresponding to the expected patterns (Supplementary Figure 7). Further improvement of prediction performances would greatly increase the applicability of the model. The required model accuracy and tolerated bias will be depend on the final context and application (e.g. research, environmental monitoring, pathogen monitoring, etc.). Aspects that can further improve model performances include increased dataset sizes for model training (Supplementary Figure 10), optimisation of acquisition settings and included fluorescence detectors (Rubbens, Props, Garcia-Timermans, *et al*., 2017) or the incorporation of different or additional stains in the cytometric measurements (Buysschaert *et al*., 2016; Duquenoy *et al*., 2020).

It should be noted that we do not expect the models to improve until the relative abundance of all taxa in a mixed community can be perfectly predicted, since flow cytometric data contain only information regarding a limited set of phenotypic properties. Studies using axenic culture data have observed that some combinations of taxa are difficult to distinguish (Rubbens, Props, Boon, *et al*., 2017; Özel Duygan *et al*., 2020), and, studies using sorting and subsequent sequencing, typically also observe sub-communities that contain multiple taxa (Zimmermann *et al*., 2016). Some taxa may be indistinguishable based on their cytometric fingerprints. Our results indicated that OTUs that are phylogenetically more closely related to each other, are more likely to be associated to the same regions in the cytometric fingerprints, and can therefore be harder to distinguish (Supplementary Figure 9). Additionally, some taxa are known to exhibit high phenotypic plasticity (Horvath *et al*., 2011), which may make it difficult for the model to reliably associate a region in the cytometric fingerprint to such taxa. This implies that we can expect that for some taxa in a given environment it may be impossible to construct performant models, despite the availability of large datasets and/or sorting data.

In contrast to previous developed methods to predict taxon abundances based on flow cytometry (Rubbens, Props, Boon, *et al*., 2017; Özel Duygan *et al*., 2020), the pipeline in our study does not rely on training models based on fingerprints of pure cultures. We have previously shown that the cytometric fingerprint of an individual taxon depends on the presence of other taxa in the community, and, that because the fingerprint of a single taxon in axenic culture and in mixed culture differs, relative abundance predictions that rely on axenic culture data may be unreliable (Heyse *et al*., 2019). Hence, the applicability of pipelines that rely on FCM fingerprints of individual taxa for model training is limited to experimental setups where it is possible to determine the *in situ* phenotypic fingerprint of individual taxa (e.g. through physical separation of cultivated taxa, cell sorting, etc.). Using cell sorting we have shown that our pipeline is able to directly link taxonomic groups to clusters in the cytometric fingerprint of both mixed and synthetic communities (Supplementary Figure 11, Supplementary Figure 8). As a result, the currently proposed pipeline is suitable for studying both environmental and synthetic communities.

### Prospects for bacterial monitoring

We used aquaculture as our model system since bacterial diseases are causing annual losses of billions of dollars worldwide in this sector (Stentiford *et al*., 2017; Shinn *et al*., 2018). These disease outbreaks are not caused by the presence of a pathogen alone, but rather by complex changes in the microbial community structure (Lemire *et al*., 2015; Dai *et al*., 2020; Huang *et al*., 2020; Infante-villamil *et al*., 2020). Additionally, the onset of mortality typically occurs very fast (Lucas *et al*., 2010; Heyse *et al*., 2021). Fast and high-throughput monitoring of bacterial community composition is a first step to mitigate the disease outbreaks, and is therefore a crucial aspect for microbial management. In practice, routine monitoring is mostly relying on (selective) plating. While these cultivation-based methods are simple and inexpensive, they remain slow (i.e. > 24h; Hallas and Monis, 2015; Rech *et al*., 2018), and provide a biased view of bacterial abundance (Van Nevel *et al*., 2017; Cheswick *et al*., 2019) and community composition (Gensberger *et al*., 2015; Sala-Comorera *et al*., 2020).

The flow cytometric toolbox for monitoring environmental communities already contains algorithms for estimating community level diversity (Props *et al*., 2016; Wanderley *et al*., 2019), stability (Liu et al., 2018) and turnover (Liu and Müller, 2020), as well as algorithms that allow to associate population dynamics with environmental or experimental parameters (Koch, Fetzer, Harms, and Müller, 2013) and pipelines that are designed for community-level classification into different categories (e.g. diseased/healthy, etc.) (Rubbens *et al*., 2020). Standalone community level metrics such as diversity or stability may be difficult to interpret, and, therefore, to couple to specific management actions, because of the high bacterial heterogeneity and fast dynamics that are typically observed in aquaculture microbiomes (Schmidt *et al*., 2017; Chun *et al*., 2018; Heyse *et al*., 2021). Additionally, different pathogens or dysbiotic states may require a different treatment. The pipeline of our study allows to add an additional layer of taxonomic information to these metrics, which will increase the actionability of the farmers. Once the models have been constructed, predictions can be made for multiple taxa simultaneously allowing to monitor a large fraction of the bacterial community.

We have shown that the pipeline that was developed in this study can be extrapolated to other applications, including analysis of laboratory mock communities and mixed reactor communities (Figure 5). In our study, average model performances on the reactor communities were lower as compared to those of the taxa in the aquaculture communities. This can be due to the smaller dataset size (i.e. 43 samples as compared to 169 for the aquaculture dataset), as this was shown to have a large influence on model performance (Supplementary Figure 10). Performances for the mock community strains was high, which can be expected due to the lower community complexity.

The main advantages of using flow cytometry for community composition monitoring lies in the speed (i.e. minutes) and the high potential for automation (Hammes *et al*., 2012; Arnoldini *et al*., 2013), which enables monitoring with high temporal resolution. Additionally, the independence of cultivation is a great advantage for monitoring managed ecosystems, since man-induced stressors, such as disinfection, are known to induce VBNC-states (Chen *et al*., 2020). Practical applications of the pipeline can include monitoring the efficacy of management strategies, follow-up disease outbreaks, monitoring the presence of probiotic strains, etc. We believe the pipeline that was developed in this study holds great potential to be integrated in routine monitoring schemes and early warning systems for biotechnological applications.

## Materials and Methods

### Samples

In this study, we used a combination of previous published flow cytometry and 16S rRNA gene amplicon data from a *L. vannamei* hatchery (Heyse et al., 2021) and new generated 16S rRNA gene amplicon data on sorted sub-communities of samples originating from this previous study. This dataset is referred to as the “aquaculture dataset”. Five gates were created for cell sorting (Supplementary Figure 1). The gates were chosen to cover the range of SYBR Green I fluorescence and side scatter that were observed in the dataset. The samples that were selected for sorting were chosen from three of the replicate tanks, over different days, in order to include communities with heterogeneous taxonomic compositions.

### Flow cytometry

Samples for flow cytometry were fixed with 5 μL glutaraldehyde (20 % vol/vol) per mL (Heyse *et al*., 2021). Glutaraldehyde-fixed, SYBR Green I-stained community samples were measured with a FACSVerse flow cytometer and sorting was performed with a BD Influx v7 Sorter USB. The procedures for flow cytometric measurements, cell sorting and control samples accompanying these procedures are outlined in detail in Supplementary Materials and Methods.

### Illumina sequencing

Sequencing of the V3-V4 region of the 16S rRNA gene amplicon sequencing was performed on an Illumina MiSeq. The DNA extraction protocols and details about the sequencing are outlined in Supplementary Materials and Methods.

### Validation datasets

The applicability of the pipeline was verified on two datasets: a synthetic community and a mixed community. The synthetic community dataset contained samples of a three strain mock community (*Stenotrophomonas rhizophila* DSM 14405, *Kocuria rhizophila* DSM 348 and *Paenibacillus polymyxa* DSM 36). The reactor community dataset originated from the study of Liu *et al*. (2019). More information regarding the validation datasets, their processing and availability is provided in Supplementary Table 2.

### Data analysis

#### Flow cytometry analysis

The flow cytometry data were imported in R (v3.6.3) (R Core Team, 2017) using the flowCore package (v1.52.1) (Hahne *et al*., 2009). The data were transformed using the arcsine hyperbolic function, and the background of the fingerprints was removed by manually creating a gate on the primary fluorescent channels (Supplementary Figure 12).

#### 16S rRNA gene amplicon sequencing analysis

Raw sequencing reads from the previous study and raw sequencing reads generated in this study were processed together. Analysis was performed with the software package MOTHUR (v.1.42.3) (Schloss *et al*., 2009). Contigs were created by merging paired-end reads based on the Phred quality score heuristic and they were aligned to the SILVA v123 database. Sequences that did not correspond to the V3–V4 region as well as sequences that contained ambiguous bases or more than 12 homopolymers, were removed. The aligned sequences were filtered and sequencing errors were removed using the pre.cluster command. UCHIME was used to removed chimeras (Edgar *et al*., 2011) and the sequences were clustered in OTUs with 97 % similarity with the *cluster.split* command (average neighbour algorithm). OTUs were subsequently classified using the SILVA v123 database. The OTU table was further analysed in R (v3.6.3) (R Core Team, 2017). OTU abundances were rescaled by calculating their proportions and multiplying them by the minimum sample size present in the data set. Absolute taxon abundances are calculated by multiplication of relative abundances with total bacterial densities as determined through flow cytometry.

#### Predictive models

##### FCM preprocessing

The data is normalized to the [0,1] interval by dividing each parameter by the maximum SYBR Green I fluorescence channel (i.e. the targeted channel) intensity value over the data set. Next, the flow cytometry data were processed by applying a Gaussian mixture mask to the data that allows to classify each cell into one of the cell clusters that are detected in the dataset. For generating the mask, all samples are subsampled to the same number of cells per sample, in order to not bias model training towards a specific sample. Similar to the method of Ludwig *et al*. (2019), the Gaussian mixture model (GMM) was optimised based on the Bayesian information criterion (BIC) using PhenoGMM (Rubbens *et al*., 2021, Supplementary Figure 13). This discretisation results in a 1D-vector for each sample that represents the number of cells present in each mixture. Unless indicated otherwise, the parameters that are included in the model are those that were optimised prior to measurement (i.e. FSC, SSC, FL1 (527/32) and FL3 (700/54)). Finally, the mixture counts were converted to relative abundances per sample and transformed using a centered log ratio (*clr*) transformation implemented in the compositions package (v. 2.0.0) (van den Boogaart and Tolosana-Delgado, 2008):

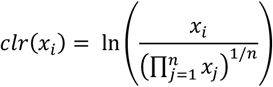

##### Illumina preprocessing

Taxa with low relative abundances are not expected to be detected through flow cytometry. Hammes *et al*., (2008) determined a quantification limit for flow cytometry of 10^2^ cells/mL. Since all samples were diluted 10 times, taxa with an absolute abundance below 10^3^ cells/mL were not expected to be observable in the flow cytometry data. Therefore, in each sample, the relative abundance of OTUs with an absolute abundance lower than 10^3^ cells/mL was set to zero.

##### Model training and validation

Models are trained for each OTU individually. To test the robustness of the pipeline, prediction performance was evaluated using independent validation sets with a nested cross-validation scheme (i.e. in the outer loop 20 % of the data is held out for validation of the final model, in the inner loop 5-fold cross-validation is used for tuning and training of the models). This outer loop was repeated three times with different fold splits. The pipeline consists of a random forest classifier to predict presence or absence of the taxon of interest and a regression ensemble (i.e. combination of a gradient boost regression and a support vector regression with polynomial kernel) to predict the relative abundance of the taxon of interest. All models were implemented using the caret (v6.0.86) (Kuhn, 2008) and caretEnsemble (v2.0.1) (Deane-Mayer, Zachary A. Knowles, 2019) packages.

Sequencing survey data is typically zero-inflated (i.e. for each individual OTU, the OTU will be absent or have a very low relative abundance; Supplementary Figure 14A). Prior to model training, samples were randomly combined *in silico* to increase the number of samples where the OTU was abundant (Supplementary Figure 14B and C). This increased model performances (Supplementary Figure 14D).

For the presence/absence classifier, samples with an OTU abundance lower than 1 % were labelled as “absent”, samples with an OTU abundance higher than 1 % were labelled as “present”. The reason why an arbitrary value of 1 % was chosen as a cut-off is that small differences in sequencing depth between samples may cause samples with similarly low relative abundances to be labelled differently (i.e. as absent or present). A random forest (RF) classifier was trained to separate both classes. Before training the classifier, the number of features was reduced using a recursive feature elimination strategy (*rfe* function in caret, 25 iterations). In short, the training data is split into a test- and trainset, the model is tuned on the train set and the features are ranked according to their importance. For each subset of the Si most important features, the model is trained on the training set and predictions are made on the test set. This procedure was repeated 25 times and the average performance profile over the different subset sizes is calculated. The performances quickly reached a plateau. To avoid incorporation of redundant features, the features required to reach an accuracy with a maximal deviation of 0.5 % of the maximal accuracy were included (*pickSizeTolerance* function in caret). Inclusion feature selection improves the ability of the model to use features/clusters that are associated to the modelled taxon, and not on correlated clusters that may belong to other taxa (Supplementary Figure 15).

For predicting the relative abundances, models with unbound outcomes were used. To avoid the generation of predictions outside the [0,1] range, the logit transformation was applied to map the relative abundances of the individual OTUs to values in the [–Inf, Inf] range before training the regression models:

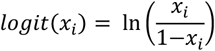

Zero values were replaced by one tenth of the smallest non-zero abundance value. The final regression predictions were inversely transformed so the final predictions were bound to the [0,1] range. A linear regression ensemble was trained using a gradient boosting regression and a support vector regression with polynomial kernel. Because the regression models were marked by a high frequency of false positive predictions, the classifier was used to correct the regression output (i.e. predicted abundances of samples for which classifier predicted “absent” were set to zero, Supplementary Figure 4).

Relative feature importance values of each model were stored to be compared either between taxa or to the sorting data. For the random forest classifier and gradient boosting regression, the mean squared error was calculated on the out-of-bag data for each tree, the values of the variable that was tested were randomly shuffled in the out-of-bag-sample and the mean squared error was calculated again. Differences in the mean squared error values were averaged and normalized. For the support vector regression, the relationship between each predictor and the outcome was evaluated by fitting a loess smoother. The R^2^ statistic was calculated for this model against the intercept only null model. This number was returned as a relative measure of variable importance.

### Data availability

The entire data-analysis pipeline is available as an R Markdown document at https://github.com/jeheyse/FCM-16S_PredictiveModelling. Raw FCM data and metadata for the aquaculture dataset are available on FlowRepository under accession ID FR-FCM-Z3CY. Raw sequence data of the bulk samples originated from a previous study (Heyse et al., 2021) and are available from the NCBI Sequence Read Archive (SRA) under accession ID PRJNA637486. Raw sequence data of the control samples, the sorted and the mock communities generated in this study are available from the NCBI Sequence Read Archive (SRA) under accession ID PRJNA691168.

## Acknowledgements

The authors would like to thank Tim Lacoere for the design of the overview figure of this manuscript, for his advice during the DNA extractions and for operating the MinION and Frederiek-Maarten Kerckhof for sharing code to analyse the MinION reads. JH is supported by the Flemish Fund for Scientific research (FWO-Vlaanderen, project 1S80618N). RP is supported by a postdoctoral fellowship of the Flemish Fund for Scientific research (FWO-Vlaanderen, project 1221020N). NB is supported by the Bijzonder Onderzoeksfonds (BOF) (BOF15/GOA/006) project. WW received funding from the Flemish Government under the “Onderzoeksprogramma Artificiële Intelligentie (AI) Vlaanderen” Programme.

## Contributions

J.H., N.B. and R.P. conceived the study. J.H. and R.P. performed the flow cytometry measurements. F.S. and S.M. performed sorting analysis. J.H. performed DNA extractions and analysed the data. R.P., P.R., W.W. advised the data-analysis. R.P. and N.B. supervised the findings of this work. J.H. wrote the paper. All authors contributed to the reviewing and editing of the manuscript. The manuscript was approved by all authors.

## Supplementary Figures

Supplementary Figure 1 – For the aquaculture dataset, cell sorting was performed on a different instrument (BD Influx v7 Sorter) as compared to the FCM measurements of community samples (BD FACSVerse). To be able to use both the community sample and the sorted sample data as a single dataset, a set of representative samples (i.e. samples originating from the replicate tanks and sampling days from which final samples for sorting were selected) was measured on both instruments and the gates that were used for sorting were manually recreated on the FACSVerse data. (A) Illustration of the five gates that were used to perform sorting on the Influx v7 Sorter. (B) Illustration of the manually recreated gates on samples that were measured on the FACSVerse. (C) Relationship between the sub-community (SC) densities in the gates drawn on data of the two instruments. The colour intensity in the first two panels is proportional to the log-scaled density of the events. Note that the colour scaling of figure A and B are independent. (Adj. R. Sq. = adjusted R-squared, C_p_ = Pearson correlation)

Supplementary Figure 2 – Community composition in samples from the aquaculture dataset. (A) Composition in the sorted samples obtained in this study. The upper title bars indicate which sub-community was sorted (i.e. “SC 1” to “SC 5”). The lower title bars indicate from which replicate tank (i.e. ‘T1” to “T5”) the community originated. (B) Composition in the non-sorted samples (data originating from the previous study, Heyse *et al*., 2021). The OTUs belonging to the 15 most abundant genera are coloured, all other genera were labelled as “Other”. The legend on the bottom applies on both panel A and B.

Supplementary Figure 3 – Number of false positive (i.e. samples incorrectly predicted to be present) and false negative (i.e. samples incorrectly predicted to be absent) samples for the classifiers that were built for the top 50 OTUs of the aquaculture dataset. Note that the number of samples are reported and not the rate. The reported p-values are the results of two sided Wilcoxon rank sum tests. The three dots for each model represent three repeated fold splits, the vertical line per OTU indicates the average performance of the replicates.

Supplementary Figure 4 – The regression ensembles frequently predicted high relative abundances for samples where an OTU was absent or present in very low abundance. To improve prediction accuracy, the predictions of the classifier were superimposed on the regression predictions: i.e. the predicted relative abundances of samples that were classified as “absent” are set to zero, predictions of samples that were classified as “present” remain unchanged. (A) Hypothetical example to illustrate the corrections that were made using the classifier predictions. Lines in blue indicate samples for which the classifier predicted “absent”, and, thus, predicted relative abundances were set to zero. Lines in white indicate samples for which the classifier predicted “present”, and, thus, the predicted relative abundance remained unchanged. (B) Illustration of predicted relative abundances for OTU 1 (*Phaeodactylibacter* sp.) from the aquaculture dataset before correction with the classifier predictions. (C) Illustration of final predicted relative abundances for OTU 1 after correction with the classifier predictions. (R. sq. = R-squared value).

Supplementary Figure 5 - Predictions for OTU2 (*Balneola* sp.; R^2^ = 0.65) from the aquaculture dataset. The five replicate shrimp cultivation tanks (“T1” to “T5”) were sampled at a resolution of 3 hours for flow cytometry and once per day for 16S rRNA gene sequencing. The presence and relative abundances for OTU2 on the time points for which no amplicon data were available were predicted in order to evaluate the ability of our approach to correctly capture dynamics of this taxon over time. The dark shades (“measured”) correspond to the values that were determined based on 16S rRNA sequencing. The lighter shades (“predicted”) correspond to time points for which only flow cytometry data was available and predictions were made using the models. Expected values can be estimated by interpolation of the measured samples (indicated with the lines between the measured samples). The reported values are averages of the two replicate measurements at each time point. (A) Predictions of the presence/absence classifier. (B) Predicted relative abundances. (C) Predicted absolute abundances, calculated by multiplying the predicted relative abundances by the total cell density as determined through flow cytometry.

Supplementary Figure 6 – Predictions for OTU6 (*Marivita* sp.; R^2^ = 0.19) from the aquaculture dataset. The five replicate shrimp cultivation tanks (“T1” to “T5”) were sampled at a resolution of 3 hours for flow cytometry and once per day for 16S rRNA gene sequencing. The presence and relative abundances for OTU6 on the time points for which no amplicon data were available were predicted in order to evaluate the ability of our approach to correctly capture dynamics of this taxon over time. The dark shades (“measured”) correspond to the values that were determined based on 16S rRNA sequencing. The lighter shades (“predicted”) correspond to time points for which only flow cytometry data was available and predictions were made using the models. Expected values can be estimated by interpolation of the measured samples (indicated with the lines between the measured samples). The reported values are averages of the two replicate measurements at each time point. (A) Predictions of the presence/absence classifier. (B) Predicted relative abundances. (C) Predicted absolute abundances, calculated by multiplying the predicted relative abundances by the total cell density as determined through flow cytometry.

Supplementary Figure 7 – Predictions for OTU13 (*Maritalea* sp.; R^2^ = 0.03) from the aquaculture dataset. The five replicate shrimp cultivation tanks (“T1” to “T5”) were sampled at a resolution of 3 hours for flow cytometry and once per day for 16S rRNA gene sequencing. The presence and relative abundances for OTU13 on the time points for which no amplicon data were available were predicted in order to evaluate the ability of our approach to correctly capture dynamics of this taxon over time. The dark shades (“measured”) correspond to the values that were determined based on 16S rRNA sequencing. The lighter shades (“predicted”) correspond to time points for which only flow cytometry data was available and predictions were made using the models. Expected values can be estimated by interpolation of the measured samples (indicated with the lines between the measured samples). The reported values are averages of the two replicate measurements at each time point. (A) Predictions of the presence/absence classifier. (B) Predicted relative abundances. (C) Predicted absolute abundances, calculated by multiplying the predicted relative abundances by the total cell density as determined through flow cytometry.

Supplementary Figure 8 – Relationship berween cluster importances assigned by the models for the top 10 OTUs in the aquaculture dataset and location of the five sorting gates in which these OTUs were detected. The colors of the dots correspond to the cluster importances that were assigned by the model. Gates in which the OTU were detected in one or more sorted sub-communities at an abundance of 1 % or higher, are indicated in blue. OTUs for which no gates are marked in blue were not found abundantly in the sorted sub-communities. The OTUs are ordered according to their R^2^ values (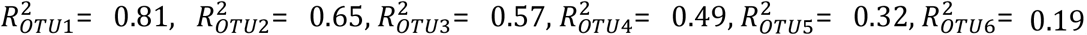, 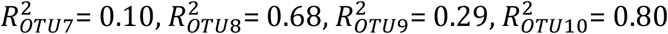).

Supplementary Figure 9 – Relationship between phylogenetic distance and similarity of model feature importances between all top 50 OTUs from the aquaculture dataset, calculated using the Bray-Curtis dissimilarities. The shaded area represents the 95 % confidence interval around the ordinary least squares regression model (p <2e-16). (Adj. R. Sq. = adjusted R-squared, C_p_ = Pearson correlation).

Supplementary Figure 10 – Learning curves to evaluate the influence of the dataset size available for training on the prediction performances for the aquaculture dataset for two OTUs: OTU1 (A & C) and OTU6 (B & D). The three dots for each model represent three repeated fold splits, the vertical line per OTU indicates the average performance of the replicates. (20 % = 34 samples, 40 % = 68 samples, 60 % = 101 samples, 80 % = 135 samples).

Supplementary Figure 11 – Correspondence of pure culture data with relative feature importance for the three strain mock community. The feature importances are averaged over the three repeats and folds. Pure culture data for *P. polymyxa* (A), *S. rhizophila* (B) and *K. rhizophila* (C). Relative cluster/feature importance of the classifier for *P. polymyxa* (D), *S. rhizophila* (E) and *K. rhizophila* (F). Relative cluster/feature importance regression ensemble for *P. polymyxa* (G), *S. rhizophila* (H) and *K. rhizophila* (I). Note that the different subplots have different colour scales.

Supplementary Figure 12 - Illustration of the cell gate applied on the inverse hyperbolic sine transformed aquaculture flow cytometry dataset. Cells are isolated from most (in-)organic and instrumental background by manual gating on the SYBR Green I fluorescence channel (533/30) and a red (> 670 nm) fluorescence channel. The colour intensity is proportional to the log-scaled density of the events.

Supplementary Figure 13 – Learning curves for the Gaussian mixture models used in this study, based on the Bayesian information criterion (BIC) (according to Rubbens *et al*., 2021). The different colours indicate different restrictions on the covariance matrices, and are indicated with a three letter code: EII (equal volumes, equal shapes, no orientation because spherical), VII (variable volumes, equal shapes, no orientation because spherical), EEI (equal volumes, equal shapes, orientation along axis), VEI (variable volumes, equal shapes, orientation along axis), EEE (equal volumes, equal shapes, equal orientation), EVE (equal volumes, variable shapes, equal orientation), VEE (variable volumes, equal shapes, equal orientation), VVE (variable volumes, variable shapes, equal orientation), EEV (equal volumes, equal shapes, variable orientation), VEV (variable volumes, equal shapes, variable orientation), EVV (equal volumes, variable shapes, variable orientation), VVV (variable volumes, variable shapes, variable orientation), EVI (equal volumes, variable shapes, orientation along axis), VVI (variable volumes, variable shapes, orientation along axis). The model with the highest BIC is retained as the final model and is indicated with the black dot. (A) For the aquaculture dataset using the scatters and two fluorescence parameter (optimum: 80 clusters, VEV). (B) For the three strain mock community (optimum: 31 clusters, VVI). (C) For the reactor communities of Liu *et al*. (2019) (optimum: 41 clusters, VEV).

Supplementary Figure 14 – (A) Relative abundance distributions of the top 50 OTUs from the aquaculture dataset, illustrating the strong zero-inflation that is typically observed in community composition survey data. (B) Distribution of the relative abundances of a random strain (OTU3) prior to the generation of *in silico* data. (C) Distribution of the relative abundances of a strain after the generation of *in silico* data. (D) Illustration of the advantage of including *in silico* generated samples for the top 3 OTUs from the aquaculture dataset. The three dots for each model represent three repeated fold splits and the vertical line indicates the average performance of the replicates.

Supplementary Figure 15 – Illustration of the added value of including a feature selection step in the pipeline for one of the taxa from the three strain mock community. (A) Pure culture data for S. rhizophila. (B) Relative cluster/feature importance for models that were trained without feature selection. (C) Relative cluster/feature importance for models that were trained with feature selection.

Supplementary Figure 16 – Relative abundances of the clusters that were detected in the microbial mock communities that were used to test for variability in flow cytometric measurements at the single-cell level, according to the recommendation of Cichocki *et al*. (2020). (A) Results for the replicates that were measured on the BD FACSVerse. (B) Results for the replicates that were measured on the BD Influx v7 Sorter USB. Note that the clusters of the two instrument are independent.

Supplementary Figure 17 – Community composition that was retrieved from the samples to evaluate the effect of glutaraldehyde in the 16S rRNA gene profile. Each test sample was sequenced in duplicate. The OTUs belonging to the 16 genera with the highest overall abundance are coloured, all other genera are labelled as “Other”.

Supplementary Figure 18 – Overview of the samples that were included to verify extraction-induced bias. (A) Community composition that was retrieved from the dilution series of the ZymoBIOMICS Microbial Community Standard (Zymo Research, USA) and the blanks, extracted with two different DNA extraction protocols (i.e. “Zymo” and “Chelex”). All contaminating OTUs are indicated as ‘Other’. (B) Sample originating from cultivation tanks that was extracted using the two DNA extraction protocols. (C) Sample originating from the algal cultures that was extracted using the two DNA extraction protocols. (D) Sample originating from the *Artemia* storage tanks that was extracted using the two DNA extraction protocols.

Supplementary Figure 19 – Overview of the samples that were included to control for potential contamination in the sorted samples. (A) Number of reads in samples from the sampling campaign (“Samples”), the buffer in which the sorted cells were collected (“Sheath”) and the Chelex solution that was used to extract DNA from the sorted samples (“Chelex”). (B) Community composition that was retrieved from the Chelex solution that was used to extract DNA from the sorted samples. One sample was taken for each of the three days DNA extractions were performed. (C) Community composition that was retrieved from the buffer in which the sorted cells were collected. One sample was taken for each day the sorting was performed. The OTUs belonging to the 16 genera with the highest overall abundance are coloured, all other genera are labelled as “Other”.

## Supplementary Tables

Supplementary Table 1 - P-values resulting from PERMANOVA analysis on the Bray-Curtis dissimilarities between the community compositions in the communities and sorted sub-communities. Note that sub-communitiy 3 was not included in the analysis since this sub-community was sorted only once. (* = For this combination it was not possible to perform PERMANOVA because the beta-dispersion of the groups was significantly differing.)

Supplementary Table 2 – Information regarding the validation datasets. Accession IDs provided for the data from the study of Liu et al., 2019 are originating from the original study. Optimisation curves for the number of clusters detected using PhenoGMM are provided in Supplementary Figure 13.

